# Single-copy Knock-In Loci for Defined Gene Expression in *C. elegans*

**DOI:** 10.1101/500892

**Authors:** Carlos G. Silva-García, Caroline Heintz, Sneha Dutta, Nicole M. Clark, Anne Lanjuin, William B. Mair

## Abstract

We have generated a single-copy knock-in loci for defined gene expression (SKI LODGE) system to insert any cDNA by CRISPR/Cas9 at defined safe harbors in the *Caenorhabditis elegans* genome. Utilizing a single crRNA guide, which also acts as a Co-CRISPR enrichment marker, any cDNA sequence can be introduced as a single integrated copy, regulated by different tissue-specific promoters. The SKI LODGE system provides a fast, economical and effective approach for generating single-copy non-silenced transgenes in *C. elegans*.

## INTRODUCTION

The *C. elegans* community has developed multiple protocols to express transgenes in this genetic model (Table 1). These protocols include extrachromosomal arrays [1], gamma/UV integration [2], biolistic transformation [3], and Mos1-mediated single copy insertion (MosSCI) [4]. The use of these tools has expedited our understanding of innumerable molecular and physiological mechanisms. However, many issues remain with these systems that limit efficacy, including inter-individual variability in expression levels, germline silencing, potential for disruption of one or more endogenous genes, laborious methodologies, and dominant phenotype co-selection markers that can influence *C. elegans* physiology (Table 1). To circumnavigate these issues we have taken advantage of precise and rapid CRISPR/Cas9 gene knock genome editing [5,6] to make a **s**ingle-copy **k**nock-**i**n **lo**ci for **d**efined **g**ene **e**xpression (SKI LODGE) system at safe harbors in the *C. elegans* genome. The SKI LODGE system allows rapid single copy tissue-specific expression of any gene. SKI LODGE uses simple PCR amplicons as repair templates along with a single well characterized targeting sequence (*dpy-10* crRNA sequence), that also facilitates Co-CRISPR selection to enrich for mutants. Furthermore, after outcrossing, SKI LODGE does not leave other alterations in the genome that may be detrimental to the organism (e.g. rescue sequences used for selection, or random insertional events that can disrupt untargeted coding or regulatory sequences), and can facilitate rapid generation of stably expressed, tissue-specific non-silenced transgenes.

**Table 1:** Characteristics of different protocols used to express transgenes in *C. elegans*.

|  | Extrachromosomal | Insertion | Bombardment | MosSCI | CRISPR | SKI LODGE |
| --- | --- | --- | --- | --- | --- | --- |
| Fast | ✓ | ✓ | ✗ | ✗ | ✓ | ✓ |
| Integrated | ✗ | ✓ | ✓ | ✓ | NA | ✓ |
| Single copy insertion | ✗ | ✗ | ✗ | ✓ | ✓ | ✓ |
| Known localization | ✗ | ✗ | ✗ | ✓ | ✓ | ✓ |
| Tissue-specific | ✓ | ✓ | ✓ | ✓ | ✗ | ✓ |
| Germline expression | ✗ | ✗ | ✓ | ✓ | ✓ | ✓ |
| Cloning-free | ✗ | ✗ | ✗ | ✗ | ✓ | ✓ |

## RESULTS AND DISCUSSION

We sought to generate transgenic *C. elegans* strains in which a single copy tissue-specific promoter had been knocked in at a defined safe harbor locus, along with the target sequence of a well characterized crRNA that could later be used to knock in any cDNA of choice by CRISPR/Cas9 (Fig. 1a). Since the discovery of genomic editing via CRISPR/Cas9, many methods have been developed to make genomic edits in *C. elegans*. One of these methods includes entirely cloning-free steps and direct injection of *in vitro*-assembled Cas9-CRISPR RNA (crRNA) trans-activating crRNA (tracrRNA) ribonucleoprotein complexes into the *C. elegans* gonad [7]. Utilizing this protocol as a base, we developed a toolkit to generate single copy insertions using only one crRNA guide. To define safe harbor loci for the SKI LODGE system, we used those that are well characterized by the MosSCI community [4] and are known to give stable expression with no silencing (Fig. 1b). We generated a suite of transgenic cassettes that have a common general design. Each SKI LODGE consists of a tissue specific promoter, followed by an epitope tag, a highly efficient CRISPR target sequence copied from the *dpy-10* gene, and a 3’ UTR for stable expression: tissue-specific promoter::3xFLAG::*dpy-10* protospacer & PAM::3’UTR (Fig. 1). By inserting 30 bases of protospacer and PAM sequence from *dpy-10* gene [8], hereafter referred to as “*dpy-10* site” we can simultaneously induce double-stranded breaks at both the transplanted SKI LODGE *dpy-10* site and the endogenous *dpy-10* locus using a single crRNA guide [9] (Fig. 2a). In addition, due to the high efficiency of the *dpy-10* site, the likelihood of getting a template inserted into the SKI LODGE cassette is amplified. In order to introduce the *dpy-10* site into the SKI LODGE cassettes, we established another easily identifiable Co-CRISPR target gene, *dpy-5* (Fig. 1a). Finally, we also added a N terminal 3xFLAG (Fig. 1a), which can be used for cell type specific biochemical applications.

**Figure 1.**
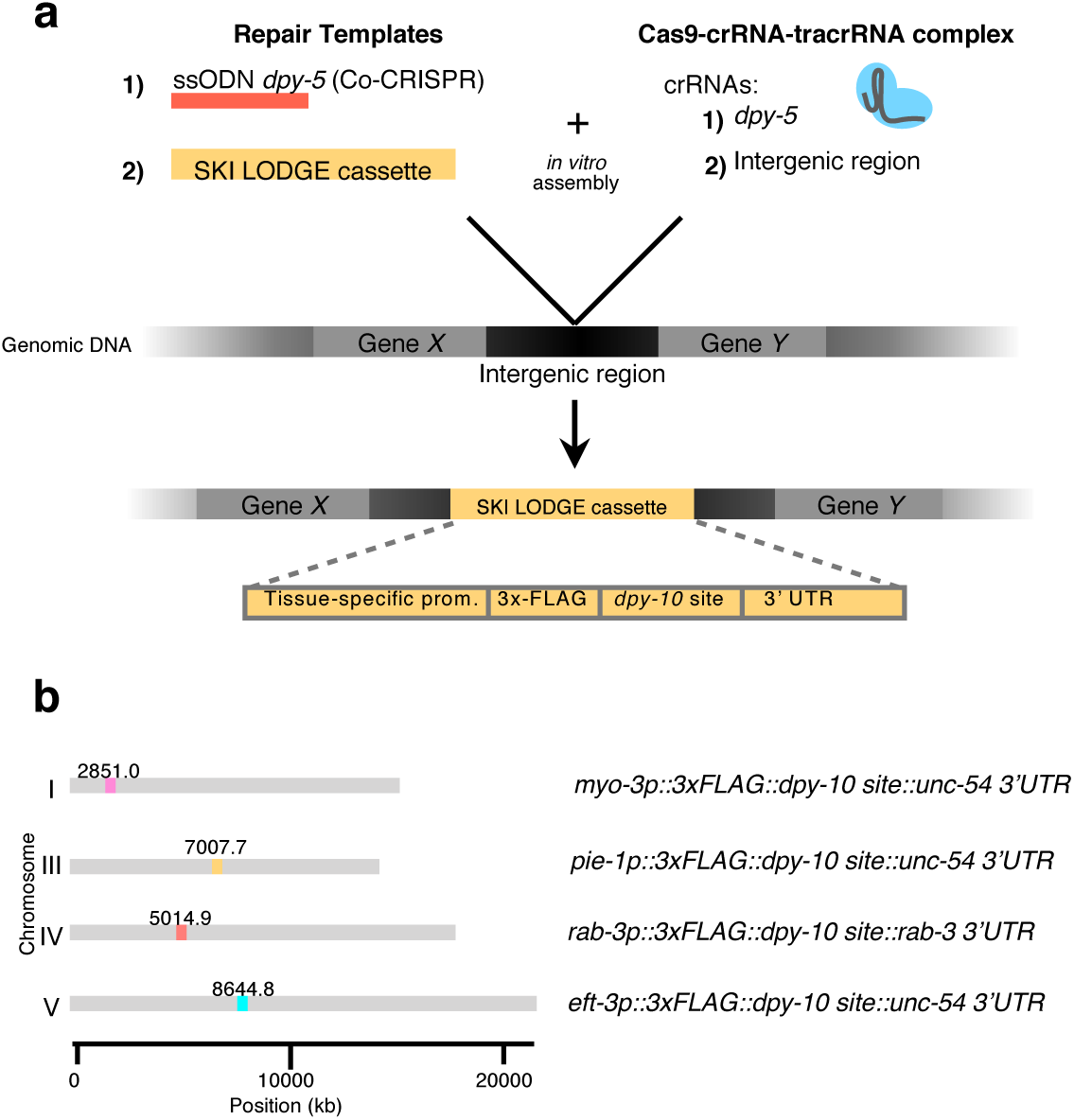
Generation of SKI LODGE lines. **(a)** Schematic of the SKI LODGE cassettes. Every SKI LODGE cassette was introduced into a defined chromosomal location. SKI LODGE PCR template(s) was combined with the CRISPR/Cas9 complex *in vitro*. This reaction mix was then injected into wild type animals. (Details about SKI LODGE construction can be accessed in Methods section). **(b)** Genomic locations of SKI LODGE insertions and composition of each cassette.

**Figure 2.**
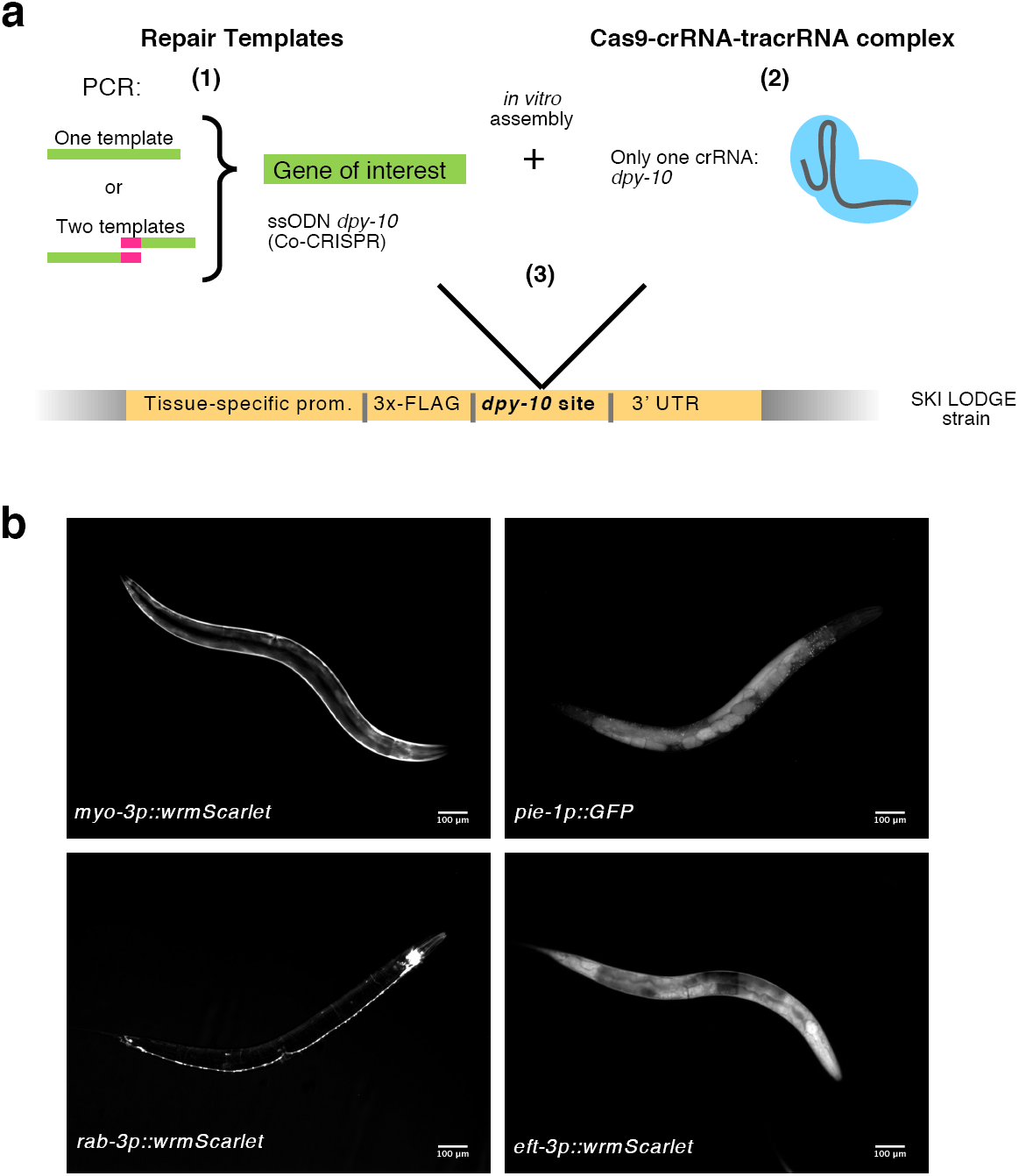
Using the SKI LODGE system to generate single-copy insertions. **(a)** Flow chart for the generation of single-copy insertions using the SKI LODGE system. (1) Select the SKI LODGE strain to express your gene of interest. Design primers to amplify your gene as homology repair template, including ∼35 bp stretches of SKI LODGE sequences immediately 5’ and 3’ to the *dpy-10* site. (2) Assemble CRISPR/Cas9 complex *in vitro*. (3) Inject pre-assembled CRISPR mix into desired SKI LODGE strain. 3-4 days post injection, isolate individual dumpy/rollers animals, and screen for desired insertion. See *step by step* section in Methods section for details. **(b)** Each SKI LODGE strain was tested by knock-in of a fluorescent protein to confirm tissue-specific expression. wrmScarlet was used to confirm ubiquitous, muscle and neuronal expression from the *eft-3, myo-3*, and *rab-3* SKI LODGE cassettes, respectively. GFP was used to confirm germline expression in the *pie-1* SKI LODGE. Animals pictured are at day 1 of adulthood.

Initially we generated four SKI LODGE strains with differing spatial expression, with each SKI LODGE cassette introduced in an intergenic region and on different chromosomes (Fig. 1b). Harboring the cassettes on different chromosomes allows the user to cross different SKI LODGE lines into each other. The SKI LODGE lines initially generated include ubiquitous (*eft-3p-eef-1A.1p-*), neuronal (*rab-3p*), muscle (*myo-3p*), and germline (*pie-1p*) promoters. We generated all strains following the Paix *et al*. protocol [7,10] (See Methods and Fig. 1a,b and Supplementary Fig. 1). After each edit, strains were outcrossed to remove any off-target and Co-CRISPR marker mutations. We modified the method of SKI LODGE cassette insertion into the *C. elegans* genome depending on the size of each promoter sequence. We did one (*rab-3, eft-3* and *pie-1*) or two (*myo-3*) edit steps to generate the final cassettes (Supplementary Fig. 1). *rab-3, eft-3* and *pie-1* were inserted using two overlapping PCR fragments, and one template was used for *myo-3*, (Supplementary Fig. 1). All final SKI LODGE lines were outcrossed a minimum of six times, and subsequently assayed for fertility, embryonic lethality, developmental timing and lifespan (Supplementary Fig. 2). Across all parameters tested, SKI LODGE strains were indistinguishable from wild type.

To verify that our SKI LODGE strains could indeed be used to drive tissue-specific gene expression, we tested all of them by CRISPR knock-in of the wrmScarlet fluorescent protein [9]. We amplified wrmScarlet gene with ∼35 bases homology arms for 3xFLAG (*myo-3, pie-1, rab-3* and *eft-3* cassettes) in the left side, and ∼35 bases homology arms for *unc-54 3’UTR* (*myo-3, pie-1* and *eft-3* cassettes) or *rab-3 3’UTR* (*rab-3* cassette) in the right side. CRISPR/Cas9 mix was assembled *in vitro [7]* using purified Cas9 protein. As predicted, utilizing only one crRNA guide (Fig. 2a), we were able to obtain *dpy-10* mutant animals that also contained inserted wrmScarlet into the SKI LODGE cassette. We observed expected patterns of tissue-specific expression for wrmScarlet driven by *eft-3, myo-3* and *rab-3* promoters (Fig. 2b and Supplementary Fig. 3). However, we did not observe expression of wrmScarlet in the germline of the *pie-1* strain (data not shown). wrmScarlet does not have introns [9], which greatly enhance transgene expression [11]. To test whether lack of intronic sequence in wrmScarlet might have impacted germline expression, we inserted GFP with artificial introns into the *pie-1* SKI LODGE. Using GFP (with intronic sequences) as a template, the *pie-1* SKI LODGE cassette drove robust GFP expression in the germline (Fig. 2b and Supplementary Fig. 3). We also observed *wrmScarlet* intestinal ectopic expression in our initial SKI LODGE single-copy *rab-3p* (neuronal) strain that contained the *unc-54 3’UTR* (data not shown). Since, 3’UTRs can modulate gene expression in *C. elegans [12]*, we generated an additional *rab-3p* SKI LODGE line, swapping out the *unc-54 3’UTR* for the *rab-3 3’UTR*. Knock-in of wrmScarlet into the *rab-3p* with the *rab-3* 3’UTR SKI LODGE line revealed no identifiable expression in the intestine at day one and six of adulthood (Fig. 2b and Supplementary Fig. 3), suggesting this line can be used to more cleanly drive gene expression in the *C. elegans* nervous system, and that use of the *unc-54* 3’UTR rather than high copy number may explain previous examples of non-neuronal leaky expression for pan-neuronal promoters such as *rab-3 [13]*. Overall, SKI LODGE strains have wild-type phenotypes with no insert, drive stable tissue-specific expression, and can be used to get DNA insertion/expression under defined promoters in as little as one week. Step by step guide for use can be found in the accompanying Methods section.

In summary, the SKI LODGE system allows insertion of single-copy transgenes in *C. elegans* into safe harbors loci using CRISPR/Cas9 editing in one week and with a high efficiency (Fig. 2a). This protocol has several advantages over existing methods (Table 1). These include, the ease of CRISPR knock-in via a highly efficient *dpy-10* crRNA guide both for knock-in and Co-CRISPR edits, and reduced time and cost due to the use of PCR amplicons and a single crRNA guide. SKI LODGE strains are phenotypically wild type, and as such are easier to inject into than mutant strains, such as *unc-119* animals often used in other methods [3,4]. The final generated transgenes are single copy, integrated at known loci that do not impact endogenous gene expression, and do not contain additional material such as selection markers or rescue constructs that impact their utility. Multiple transgenic lines generated using the same SKI LODGES ensures comparable levels of expression, identical regulatory sequences, no position effects, and the same genetic background. SKI LODGE also allows tissue specific epitope tagging for future biochemical applications such as IP, ChIP, IF, or Western blotting. SKI LODGE lines can be used to insert one or two templates (with overlapping sequences) at the same time (Fig. 2a). However, for large insertions (>3000 bp) we recommend following protocols that use plasmid templates with long homology arms [14]. We will continue to develop new SKI LODGE lines with enhanced application (see Supplementary Table 1 for pipeline) and make them available freely to the community.

## MATERIALS AND METHODS

### Step by step guide to insert your favorite gene into the SKI LODGE strains

#### Step 1

Select the SKI LODGE strain to express your gene of interest

#### Step 2

Design primers with homologous recombination arms that contain at least 35 bp of SKI LODGE 3xFLAG sequences at the 5’ end of the Forward primer, and at least 35 bp of the SKI LODGE 3’UTR sequences at the 5’ end of the Reverse primer. Use the following model and table as a reference for primer designing.

Note: In your primer design, make sure the transcript to be inserted is in frame with the SKI LODGE epitope tag. The final correct edit should have your gene of interest in frame with the N-terminal epitope tag, followed immediately by SKI LODGE 3’ UTR sequences. The entire *dpy-10* site will be removed.

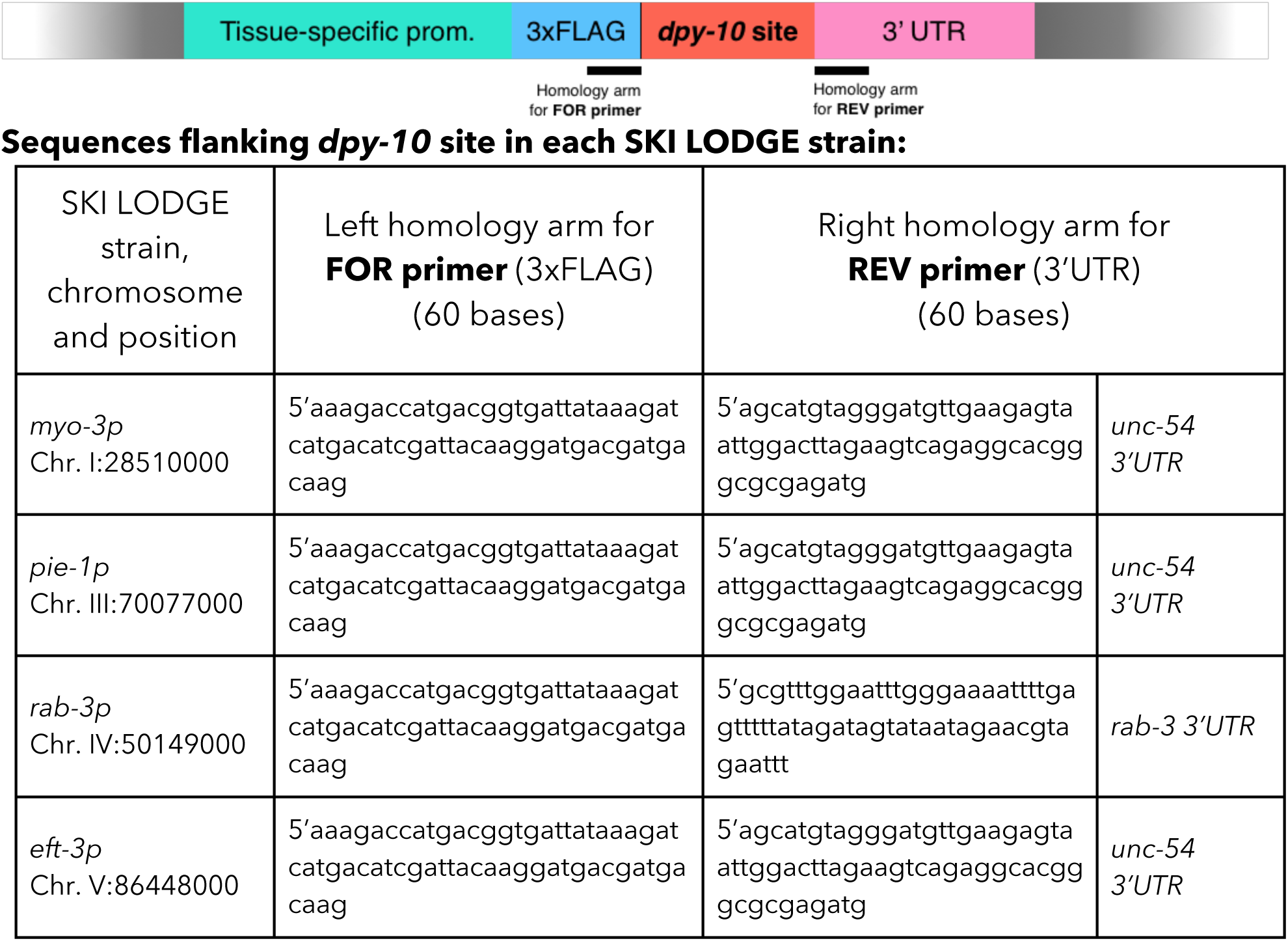

#### Step 3

Make a mastermix with your newly designed primers and amplify your homology repair template by PCR using the following conditions:

| Reagent | Volume (μl) |
| --- | --- |
| H<sub>2</sub>O | Up to 300 |
| FOR primer (100 μM) | 1.5 |
| REV primer (100 μM) | 1.5 |
| DNA template | 400-600 ng |
| Phusion 2x Master Mix | 150 |

Divide mastermix between 6 PCR tubes, 50 µl per tube. Use the following gradient and PCR cycle conditions:

| Tube | Tm | PCR cycle steps |  |  |
| --- | --- | --- | --- | --- |
| 1 | 60 °C | **1** | 98° 2' |  |
| 2 | 62 °C | **2** | 98° 30'' | 35x<br>cycles |
| 3 | 64 °C | **3** | **Gradient 30''** |  |
| 4 | 66 °C | **4** | 72° * |  |
| 5 | 68 °C | **5** | 72° 10' |  |
| 6 | 70 °C | **6** | 4° ∞ |  |

\*Extension time will depend of the length of your gene of interest. The typical recommendation is to allow 1 minute of extension time for every 1 kb to be amplified.

#### Step 4

Check PCR products on a gel to verify that the Homologous Recombination (HR) template has amplified as a clean single band. Add 5 µl of 10x orange loading buffer to each sample and run 5 µl on a 1% agarose gel.

#### Step 5

Purify and concentrate your homology repair template by pooling all reactions that amplified as a strong single band into a single sample, and pass it through a Qiagen minelute column (#28006). Follow kit instructions, and elute with 10 µl of water. Determine the concentration of your purified template.

#### Step 6

Assemble of CRISPR/Cas9 complex *in vitro*:

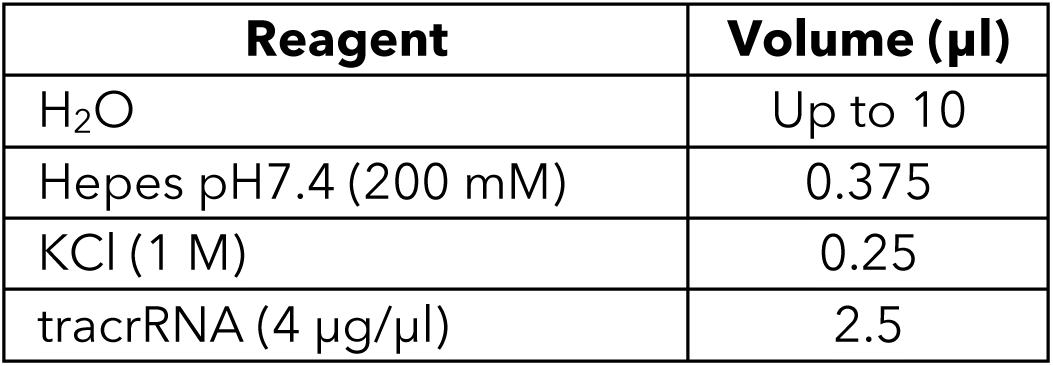

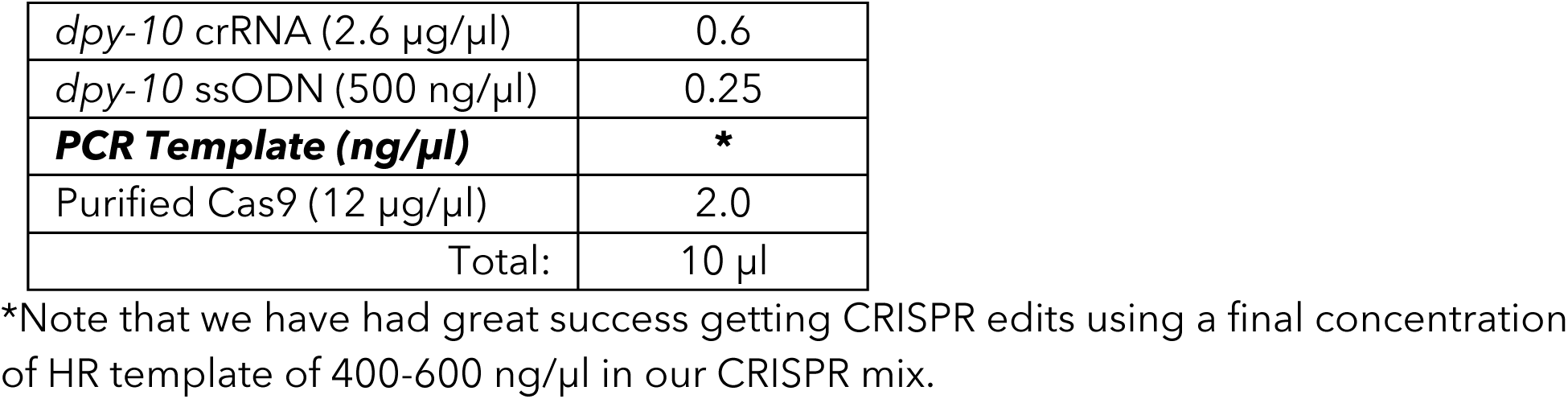

Injection mixes can be prepared in advance without Cas9 protein, separated into 2 tubes of 4 μl, and stored at −80°C. We have found that frozen mixes stored up to one year at −80°C are still effective at generating CRISPR edits. Before injecting, thaw one 4 μl mix, add 1 μl purified Cas9 (12 μg/μl), mix by pipetting, spin for 2 min at 13000 rpm, and incubate at 37°C for 10 minutes. Inject into day 1 adults of the relevant SKI LODGE line.

#### Step 7

3-4 days after injection, screen for plates that have produced many dumpy and/or roller progeny. From these plates, individually plate single *dpy* animals and allow them to lay eggs before screening them for your desired edit, either by looking for fluorescent expression and/or genotyping them by PCR. We have annotated forward primers close to the end of every tissue-specific promoter, and reverse primers at the beginning of 3’UTR that can be used for screening, genotyping, and sequencing your desired edit. Use the next figure and table as a reference for primer designing. Note: if you design new primers for genotyping, choose them outside of the homology arms to look for successful insertion.

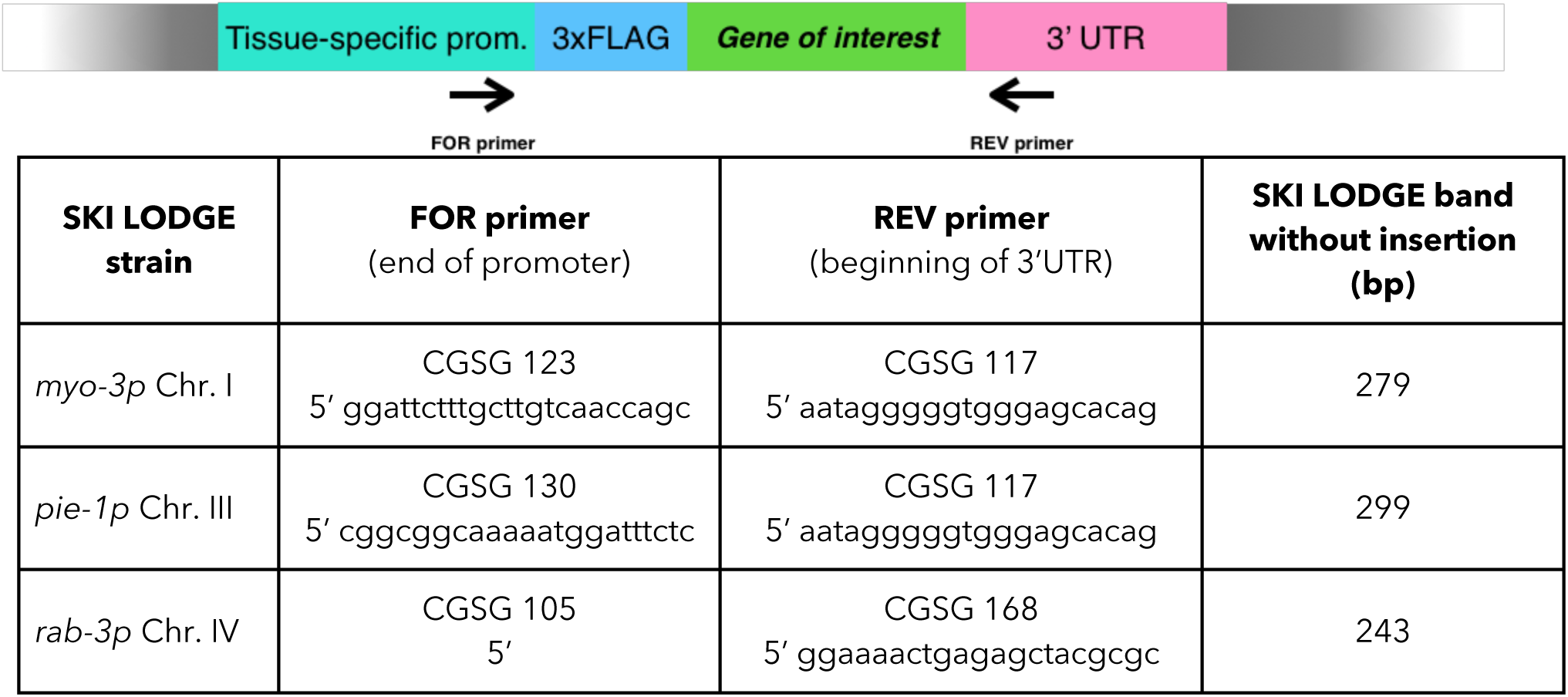
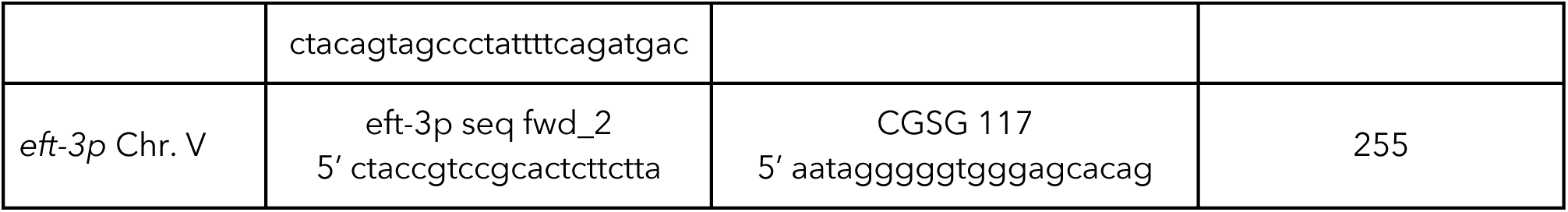

#### Step 8

Confirm that the knock-in CRISPR edit is correct by sequencing.

#### Step 9

Outcross your newly made SKI LODGE line to get rid of the *dpy-10* Co-CRISPR edit, as well as any other off-site mutations. Note that a good outcrossing genotyping strategy is necessary since outcrossing will be done using the N2 strain and not SKI LODGE strains.

The following figure and table show primers flanking the SKI LODGE cassette that can be used to outcross the new SKI LODGE line to N2. We also recommend designing an internal Forward primer close to the 3’ end of your knocked in gene. Using this 3 primer PCR strategy, you will obtain a SKI LODGE knock-in band around 1000 bp.

Figure of final SKI LODGE strain after knock in, and positions for primers:

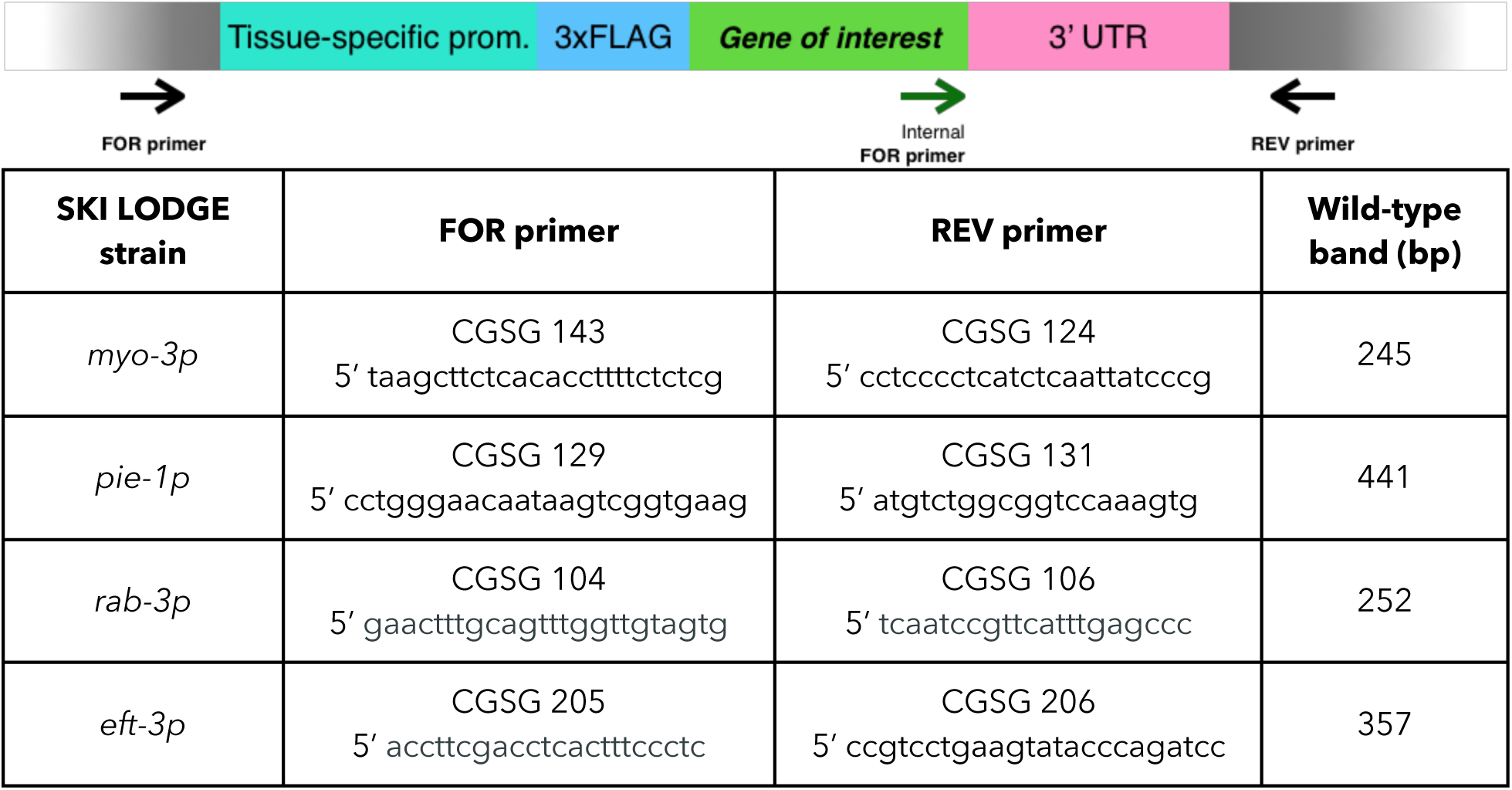

### Strains and maintenance

Bristol N2 strain was provided by the Caenorhabditis Genetics Center (CGC), University of Minnesota. All worms were grown on nematode growth medium (NGM) agar plates seeded with the *Escherichia coli* bacterial strain OP50-1 and maintained using standard procedures [15].

### Strains generated in this work and sequences

All strains generated in this work are listed in Table 4, and will be available for distribution through the CGC. All sequences files for SKI LODGE lines are available on request. All oligonucleotides, crRNAs, tracrRNA and repair templates used in this study are listed in Table 2 and 3.

**Table 2:**
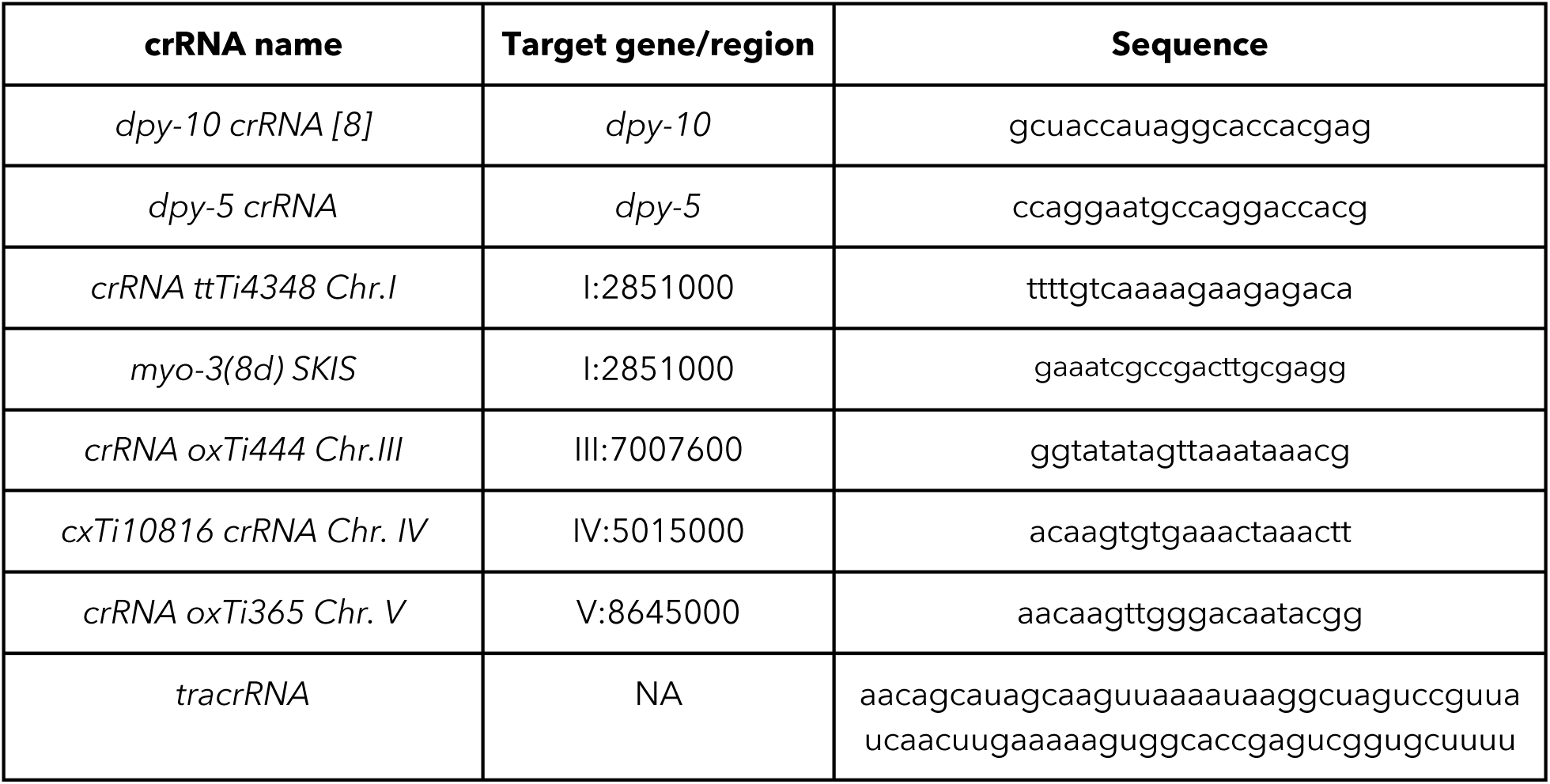
crRNAs and tracrRNA used in this work.

**Table 3:**
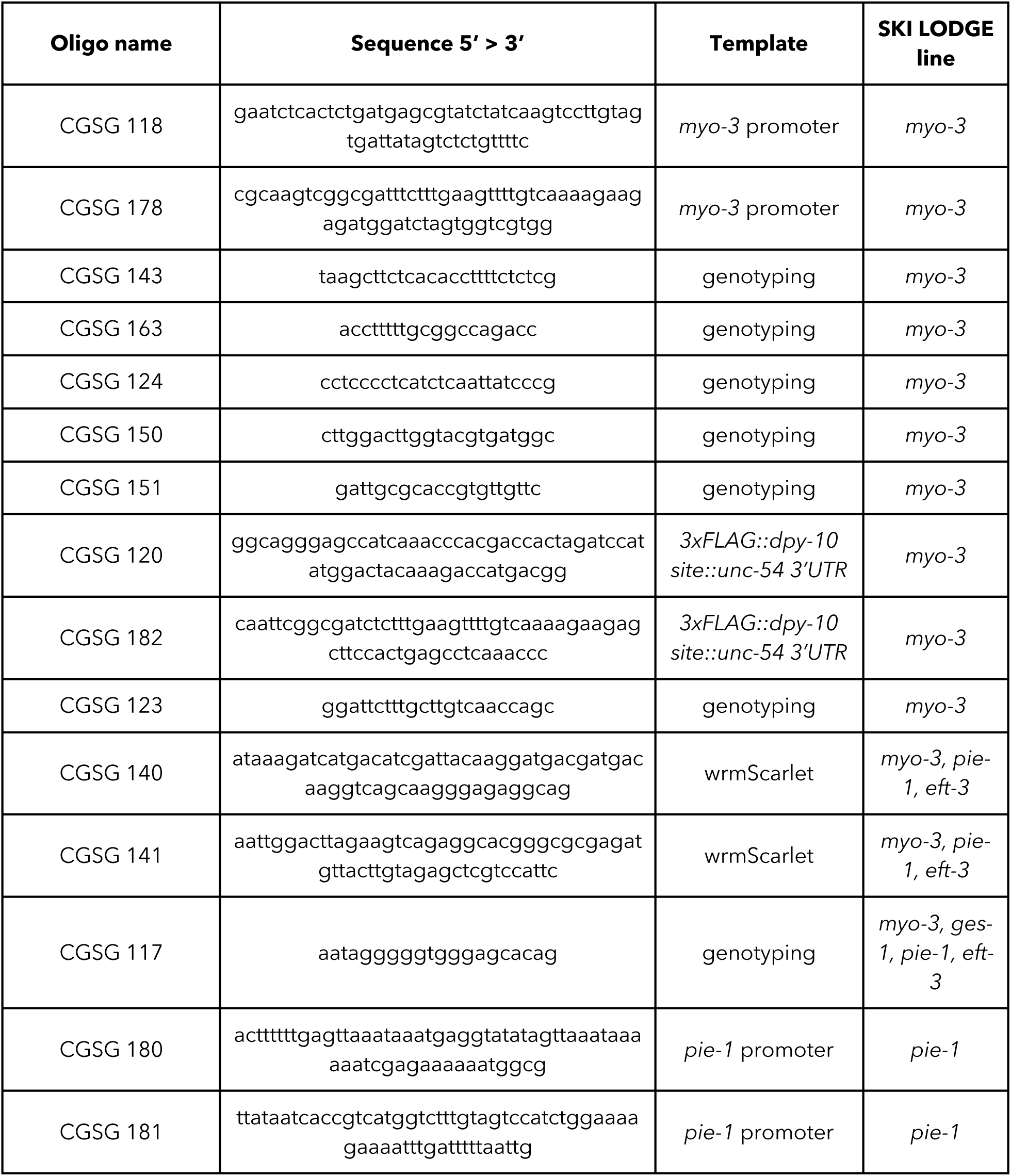
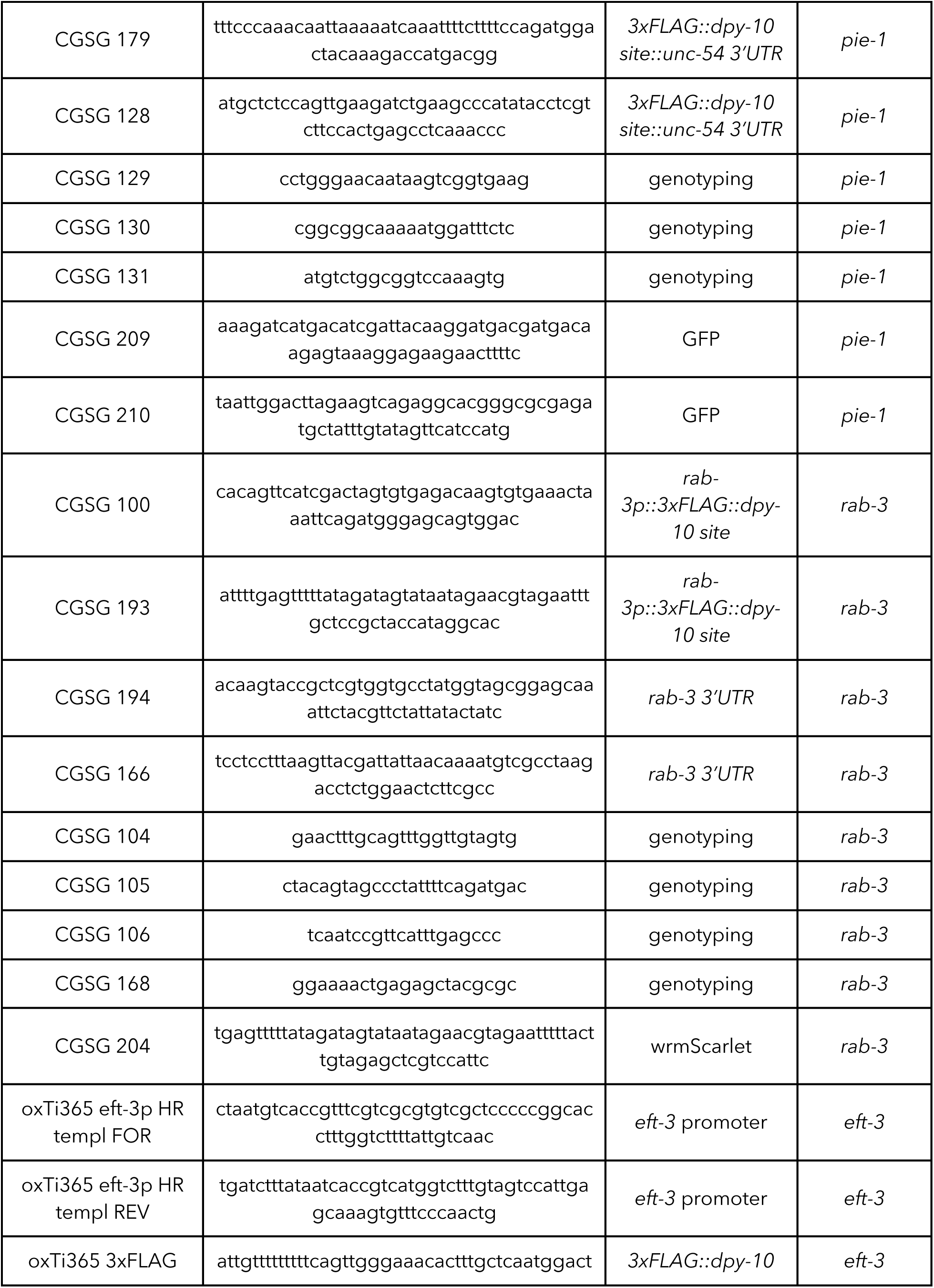
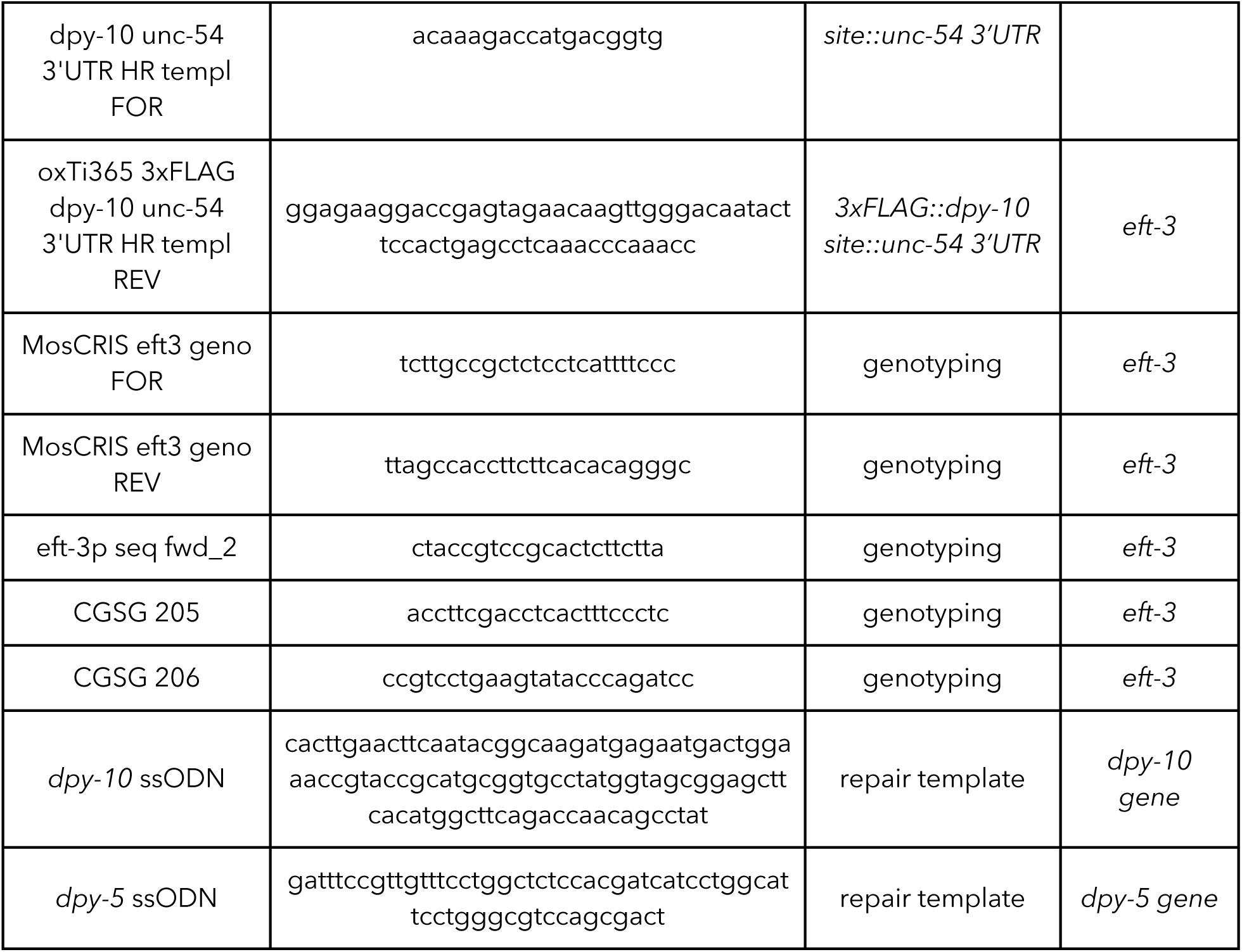
Oligonucleotides used in this work to generate SKI LODGE strains.

**Table 4:** Strains generated in this work.

| Strain | Genotype | crRNA used | Recipient strain |
| --- | --- | --- | --- |
| **SKI LODGE strains** |  |  |  |
| WBM1119 | *N2, wbmls60[pie-1p::3XFLAG::dpy-10 crRNA::unc-54 3'UTR, III:7007600]* | crRNA oxTi444 Chr.III | N2 |
| WBM1126 | *N2, wbmls61[myo-3p::3XFLAG::dpy-10 crRNA::unc-54 3'UTR, I:2851000]* | crRNA ttTi4348 Chr.I<br>*myo-3(8d)* SKIS | N2 |
| WBM1140 | *N2, wbmls65[eft-3p::3XFLAG::dpy-10 crRNA::unc-54 3'UTR, V:8645000]* | crRNA oxTi365 Chr. V | N2 |
| WBM1141 | *N2, wbmls66[rab-3p::3XFLAG::dpy-10 crRNA::rab-3 3'UTR, IV:5015000]* | cxTi10816 crRNA Chr. IV | N2 |
| WBM1163 | *N2, wbmls76[eft-3p::3XFLAG::dpy-10 crRNA::unc-54 3'UTR, IV:5015000]* | *cxTi10816 crRNA Chr. IV* | N2 |
| **Tested SKI LODGE strains** |  |  |  |
| WBM1133 | *N2, wbmls63[myo-3p::3XFLAG::wrmScarlet::unc-54 3'UTR, \*wbmls61]* | *dpy-10* | WBM1126 |
| WBM1143 | *N2, wbmls67[eft-3p::3XFLAG::wrmScarlet::unc-54 3'UTR, \*wbmls65]* | *dpy-10* | WBM1140 |
| WBM1144 | *N2, wbmls68[rab-3p::3XFLAG::wrmScarlet::rab-3 3'UTR, \*wbmls66]* | *dpy-10* | WBM1141 |
| WBM1153 | *N2, wbmls72[pie-1p::3XFLAG::GFP::unc-54 3'UTR, \*wbmls60]* | *dpy-10* | WBM1119 |
| **Other strains generated in this work** |  |  |  |
| WBM1043 | *N2, wbmls53[rab-3p::3XFLAG::dpy-10 crRNA::unc-54 3'UTR, IV:5015000]* | *cxTi10816 crRNA Chr. IV* | N2 |
| WBM1063 | *N2, wbmls57[rab-3p::3XFLAG::wrmScarlet::unc-54 3'UTR \*wbmls53]* | *dpy-10* | WBM1043 |
| WBM1139 | *N2, wbmls64[pie-1p::3XFLAG::wrmScarlet::unc-54 3'UTR \*wbmls60]* | *dpy-10* | WBM1119 |
| WBM1164 | *N2, wbmls77[eft-3p::3XFLAG::GFP::unc-54 3'UTR, \*wbmls65]* | *dpy-10* | WBM1140 |
| WBM1165 | *N2, wbmls78[eft-3p::3XFLAG::GFP::unc-54 3'UTR, \*wbmls76]* | *dpy-10* | WBM1163 |

### Identification of harbor loci

To identify safe harbor loci to knock in the SKI LODGE cassettes, we analyzed genomic regions used in the MosSCi system [4] (Fig. 1b). SKI LODGE sites were chosen using the following basic requirements: each was in an intergenic region so as not to affect other gene expression or function, and each had a strong crRNA guide site as determined by MIT CRISPR design (http://crispr.mit.edu) and/or Benchling (https://benchling.com) platforms (Table 2). We chose the following chromosome positions to insert SKI LODGE cassettes: *myo-3p* (Chr. I:28510000), *pie-1p* (Chr. III:70077000), *rab-3p* (Chr. IV:50149000) and *eft-3p* (Chr. V:86448000).

### Cas9 purification

Cas9-His-tagged was purified by overexpressing in *E. coli* BL21 Rosetta cells. Nickel-NTA beads were used to pull it down followed by HPLC purification. Briefly, we inoculated LB culture with SpCas9 containing *E. coli* Rosetta cells and shook for 3-6 h until the culture reached an OD of 0.6-0.8. The culture was cooled down on ice water to below 20°C, and 0.5 mM IPTG was added followed by incubation overnight in a shaker at 20°C. The culture was spun, and the bacterial pellet was resuspended in 27.5 ml of 50 mM Tris pH 8, 150 mM NaCl, 10% glycerol and 2 mM TCEP. Cells were lysed by adding 2 ml 10X FastBreak buffer (Promega V8571) and 5 μl Benzonase, incubated for 15-20 minutes at RT, and spun at 38000g for 15 minutes at 4°C and the supernatant collected. Ni-NTA resin was washed in an equilibration buffer (50 mM Tris pH 8, 500 mM NaCl and 10% glycerol). The supernatant was then incubated with 1 ml of this pre-washed Ni-NTA resin at 4°C for 1 h. The sample was poured into a disposable column and washed with wash buffer (20 ml of 50 mM Tris pH 8, 500 mM NaCl and 10% glycerol, 20 mM Imidazole and 2 mM TCEP). The sample was then eluted using elution buffer containing 5 ml 50 mM Tris pH 8, 500 mM NaCl and 10% glycerol, 400 mM Imidazole and 2 mM TCEP. The eluted Cas9 protein was diluted 1-2 fold with 1X PBS and loaded onto a 5 ml heparin column equilibrated with 1X PBS. The sample was eluted in a linear salt gradient from 0.1 to 1 M NaCl in 1X PBS and 1 ml fractions were collected. Fractions were analyzed on a gel and pooled all Cas9 containing fractions followed by concentrating them down to 3 ml volume with Amicon Ultra-15 filter (30 kDa cutoff, spun 4000 g 20 minutes at 4 °C). We exchanged the buffer to 1X PBS, 10% glycerol and 2 mM TCEP using a PD-10 column. The insoluble protein was spun down at 4°C for 15 min and concentrated down to a final volume of 500 μl. Purified Cas9 was filter sterilized (0.22 μm) and the final concentration was measured by Nanodrop and stored at −80°C.

### Microinjection and CRISPR/Cas9-triggered homologous recombination

All CRISPR edits and insertions required to generate the SKI LODGE strains were performed using the CRISPR protocol developed by [7]. Briefly, homology repair templates were amplified by PCR, using primers that introduced a minimum stretch of 35 bp homology to the SKI LODGE cassette at both ends. (For more information please see the step-by-step guide to follow). CRISPR injection mix reagents were added in the following order: 0.375 µl Hepes pH 7.4 (200mM), 0.25 µl KCl (1M), 2.5 µl tracrRNA (4 µg/µl), 0.6 µl *dpy-10* crRNA (2.6 μg/μl) or 0.5 µl *dpy-5* crRNA (8 μg/μl), 0.25 μl *dpy-10* ssODN (500 ng/μl) or 0.5 μl *dpy-5* ssODN (1000 ng/μl), and PCR repair template(s) up to 500 ng/µl final in the mix. Water was added to reach a final volume of 8 µl. 2 µl purified Cas9 (12 μg/μl) added at the end, mixed by pipetting, spun for 2 min at 13000 rpm and incubated at 37°C for 10 minutes. Mixes were microinjected into the germ line of day 1 adult hermaphrodite worms using standard methods [2].

### SKI LODGE cassettes construction

All strains were generated by CRISPR protocol described above using ∼35 bp of homology as recombination arms (Supplementary Fig. 1, and see Table 2 for a list of all primer sequences used for generate homologous recombination templates by PCR). In order to introduce the *dpy-10* site into the SKI LODGE cassettes, we established another easily identifiable Co-CRISPR target gene, *dpy-5*. *dpy-5* gene has only one exon and a well-defined dumpy phenotype [15]. After each edit, the modified strains were outcrossed to remove the Co-CRISPR marker mutation (*dpy-10* or *dpy-5*), as well as any additional unwanted off-site mutations. To generate the *myo-3* SKI LODGE on chromosome I:28510000 (Supplementary Fig. 1), 2573 bp of *myo-3* promoter was introduced, using *dpy-10* as a Co-CRISPR marker. Then, 806 bp of *3xFLAG::dpy-10 site::unc-54 3’UTR* cassette was added, using *dpy-5* as a Co-CRISPR marker. The *pie-1* SKI LODGE on chromosome III:70077000, the *rab-3* SKI LODGE on chromosome IV:50149000, and *eft-3* SKI LODGE on chromosome V:86448000 were each made using two HR repair templates with overlapping sequence that were co-injected as part of the same CRISPR mix, using *dpy-5* as Co-CRISPR marker (Supplementary Fig. 1). The templates to make or the *pie-1* cassette were 1166 bp of *pie-1* promoter and 806 bp of *3xFLAG::dpy-10 site::unc-54 3’UTR*. The templates to make the *rab-3* cassette, were 1405 bp of *rab-3p::3xFLAG::dpy-10 site* and 732 bp of *rab-3* 3’ UTR. The templates to make the *eft-3* cassette were 682 bp of the *eft-3* promoter and 836 bp of *3xFLAG::dpy-10 site::unc-54 3’UTR*. All final SKI LODGE strains were sequence verified and outcrossed at least six times to N2.

### Screening in worms with dumpy phenotype and genotyping

Immediately after injection, individual worms were placed at 20°C. 3-4 days after injection, all plates were screened for progeny that contained the relevant Co-CRISPR edit. ∼150 F1 dumpy/roller animals were singled out, one per plate, to lay eggs. After 2 days, F1 worms were placed in 5 µl of single worm lysis buffer (30 mM Tris pH 8.0, 8 mM EDTA pH 8, 100 mM NaCl, 0.7% NP-40, 0.7% Tween-20 and 100 µg/ml proteinase K) and lysed for 1 h at 60 °C, followed by incubation at 95 °C to inactivate the proteinase K. We then screened for the CRISPR edit by PCR using Apex Taq RED Master Mix 2.0X (Genesee Scientific) as recommended by the manufacturer, using 1 µl of worm lysate as a template (see Table 3 for a list of all primers used for PCR genotyping). F2 progeny from those plates that genotyped positive for the desired edit were again genotyped and used to amplify the full cassette. PCR amplicons were then purified using the QIAquick PCR purification kit (Qiagen) as recommended by the manufacturer. Purified PCR products were then sequenced by Sanger sequencing methods (Genewiz).

### Verification of SKI LODGE expression

SKI LODGE strains were tested by knock-in of 696 bp of wrmScarlet amplified from pSEN89 and or by knock-in of 867 bp of eGFP amplified from pIM26. A minimum of 35 bp were used as homologous recombination arms.

### Analysis of phenotypes

To study the fertility of SKI LODGE strains, seven hermaphrodites were individually selected at the L4 larval stage and transferred to new OP50-1 plates every 24 h over the course of 6 days at 20°C. Plates were scored for dead embryos and viable offspring. Embryos that did not hatch within 24 h after being laid were considered dead. To analyze developmental timing, animals were synchronized by bleaching. After bleaching, embryos were incubated overnight at 20° C in 1X M9 buffer and gentle rocking to allow hatching. After overnight incubation, OP50-1 plates were seeded with L1 larvae. Developmental stages were evaluated every 24 hours under a dissecting scope and scored for frequency of each developmental stage present.

Lifespan experiments were performed as described previously [16]. Before the start of each lifespan experiment, gravid adult worms were bleached and eggs were seeded on HT115 bacteria (100 eggs per plate) at 20° C until they reached adulthood (72 hr), at which point 100 adult worms per treatment were transferred to fresh plates with 20 worms per plate. Worms were transferred to fresh plates every other day until reproduction had ceased (day 10-12). Survival was scored every 1–2 days and a worm was deemed dead when unresponsive to 3 taps on the head and tail. Worms were censored due to contamination on the plate, if they crawled up onto the walls, had eggs hatching inside, or exhibited pronounced vulval protrusion.

### Microscopy

DIC and fluorescence imaging of whole worms was performed using a Zeiss Discovery V8 and Apotome.2-equipped Imager M2 microscope equipped with Axiocam cameras. Worms were anesthetized in 40 mM Sodium Azide diluted in 1X M9 and mounted on 2% agarose pads on glass slides.

### Statistical analysis

GraphPad Prism was used for statistical analyses. Mann-Whitney test was used to analyze fertility and embryonic lethality. The Log-rank (Mantel-Cox) analysis was used to compare survival curves: n = 85-100 worms. For all experiments p values < 0.05 were considered significant.

### Ethics statement

*C. elegans* is not protected under most animal research legislation.

## ACKNOWLEDGEMENTS

We thank Henning Arlt for his generous help and expertise in the purification of Cas9 protein. We thank the Caenorhabditis Genetics Center for providing the N2 strain. We also thank members of the Mair laboratory for helpful discussions and for comments on the manuscript. C.G.S-G. is funded by the Yerby Postdoctoral Research Fellowship. CH is funded by the Charles A. King Trust Postdoctoral Research Fellowship. WBM is funded by: NIH R01AG044346, NIH RO1AG059595, NIH R21AG056930 and NIH R01AG054201.

## SUPPORTING INFORMATION

**Supplementary Figure 1.**
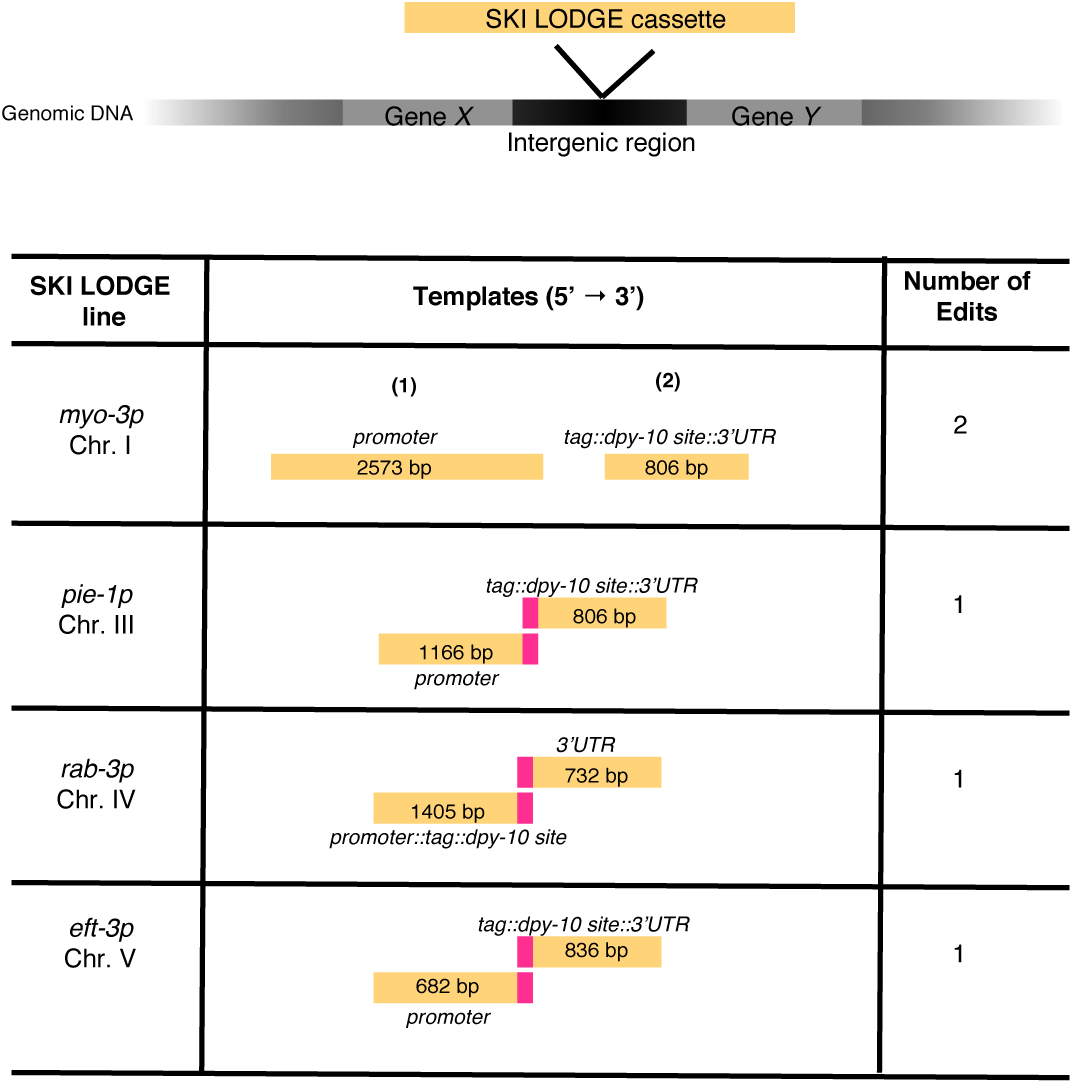
Templates used to generate SKI LODGE lines. SKI LODGE cassettes were introduced into intergenic regions on different chromosomes. Table shows templates that were used to insert SKI LODGE cassettes, and the number of CRISPR/Cas9 edits required to make the final line.

**Supplementary Figure 2.**
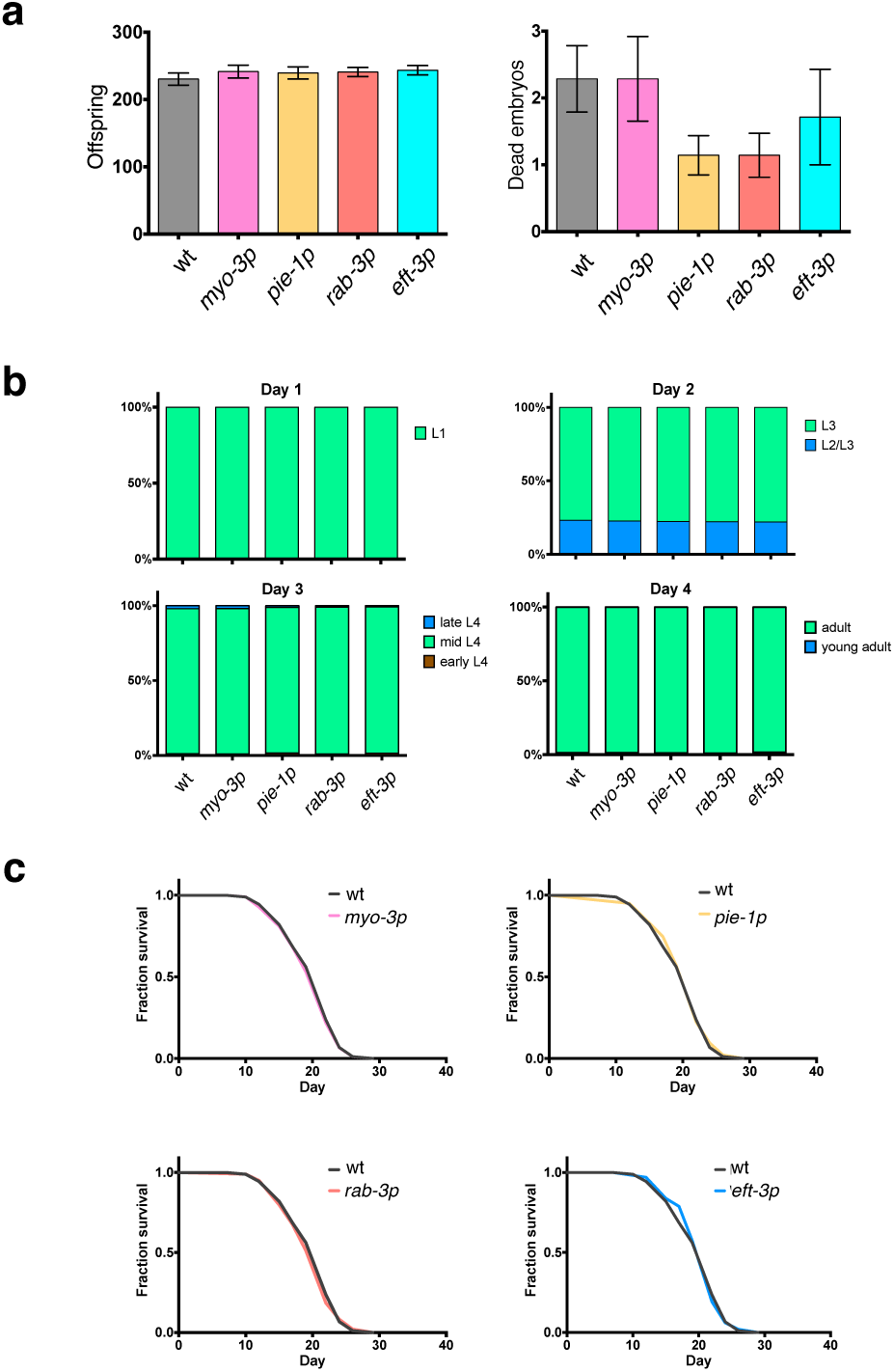
Phenotypes analyzed in SKI LODGE lines. All SKI LODGE lines produced wild type numbers of progeny and wild type levels of embryonic lethality **(a)**, as well as wild type developmental timing **(b)** and lifespan **(c)**. Values shown in a and b are an average of two independent replicates, error bars represent SEM. c is showing one replicate of two.

**Supplementary Figure 3.**
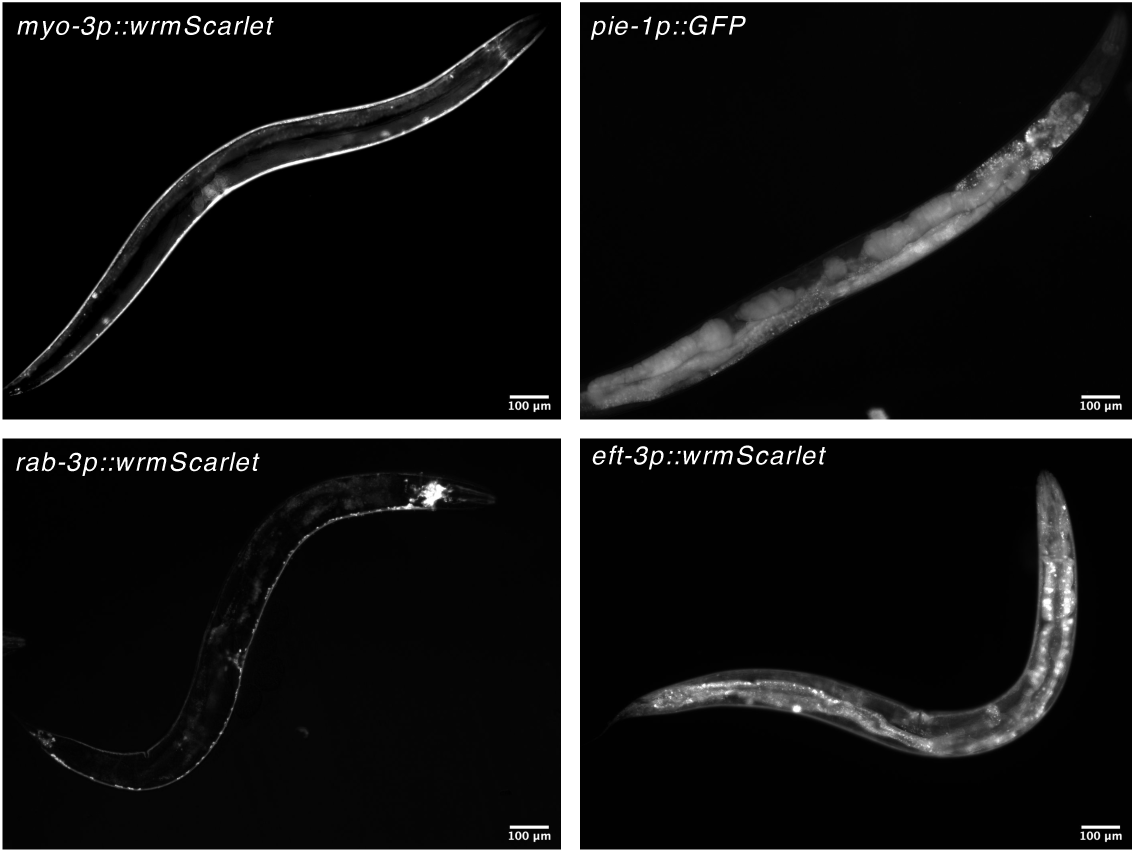
SKI LODGE lines maintain tissue-specific expression with age. SKI LODGE strains were tested by knock-in of a fluorescent protein. wrmScarlet was used to confirm ubiquitous, muscle and neuronal expression from the *eft-3, myo-3*, and *rab-3* SKI LODGE cassettes, respectively. GFP was used to confirm germline expression in the *pie-1* SKI LODGE. Pictures of six-day old animals.

**Supplementary Table 1:** Future pipeline for SKI LODGE lines.

| Aim | Description |
| --- | --- |
| Add additional promoters | We will add additional tissue-specific promoters. |
| Rapid promoter switching for any desired promoter | Flank SKI LODGE promoters with matching crRNA site sequence, enabling one step promoter switching. |
| Introduce SL2 sequence | Introduce an SL2 sequence after the *dpy-10* site to facilitate trans splicing with a fluorescent protein without generating fusions. |
| Fluorescence selection | Modified SKI LODGE cassette with *dpy-10::STOP::fluorescent protein*. Knock in will remove *dpy-10::STOP* to allow selection by fluorescence |

